# Whole *Aegilops tauschii* transcriptome investigation revealed nine novel miRNAs involved in stress response

**DOI:** 10.1101/525972

**Authors:** Behnam Bakhshi, Ehsan Mohseni Fard

## Abstract

*Aegilops tauschii* is wild relative of bread wheat. This species has been reported as donor of bread wheat D genome. There are also many reports that mentioned *Ae. tauschii* importance in biotic and abiotic stress resistance. Thus, it is important to investigate regulatory mechanisms involved in *Ae. tauschii* stress tolerance. MiRNAs have been reported as essential regulatory elements in stress response. In this study, using ESTs/TSAs databases nine novel miRNA stem-loops have been identified including ata-miRNovel_1 to 9. MiR156, miR168, miR169, miR319, miR397 and miR530 are some miRNA families that their mature miRNAs could be matched with these novel stem-loop miRNAs. We have also observed wheat ESTs/TSAs as similar as *Ae. tauschii* ESTs/TSAs that could produce new miRNA stem-loops in wheat. These novel miRNA stem-loops could play important role in transcription regulation involved in stress response.

## Introduction

*Ae. tauschii* is a wild relative species of bread wheat with wide distribution in the Mediterranean and western Asia. This diploid species is included with D genomes that composed of 14 chromosomes ^1-3^. *Ae. tauschii* and bread wheat (*T. aestivum*) are in common in D genome. Thus, *Ae. tauschii* species is considerably important for transferring appropriate traits to bread wheat and also increasing genetic diversity of bread wheat. This species has been introduced as one of the most important genetic resources for bread wheat breeding programs especially for improving bread wheat in stress response ^4-8^. Therefore, it is important to identify regulatory mechanisms involved in *Ae. tauschii* against stress response and also application of resources in bread wheat breeding program. MiRNAs are essential regulatory elements in stress response regulatory mechanisms ^9^. MiRNAs are small 19-24 nt regulatory RNAs that are encoded by endogenous MIRNA genes ^9^. MiRNA biogenesis includes three steps. At first, RNA Polymerase II produces primary miRNA transcript (pri-miRNA) from MIRNA genes. The next step is processing of pri-miRNA to precursor miRNA (pre-miRNA) completed by Dicer like one (DCL1) with the association of proteins involved in the biogenesis of miRNAs like HYL1 and SE. Pre-miRNA has a stem-loop structure that reform to mature miRNA via further processing by DCL1 ^9^. This mature miRNA could inhibit genes expression by either cleavage or translational inhibition of mRNAs ^10^. MiRNAs involved in plant developments and physiology as well as phytohormones ^11^. These features put miRNAs at the center of regulatory expression networks ^12^.

Significant roles of miRNAs made researchers to look up intensively for novel miRNAs. To this end, expressed sequence tags have been used widely ^13-16^. Pani and Mahapatra have predicted two new miRNAs in a medicinal plant *C. roseus*. It has been observed that these miRNAs are involved in vinblastine and vincristine biosynthesis ^15^. Furthermore, computational prediction has been also used to detect potential miRNAs in *H. annuus*. This prediction led to identify three potential miRNAs that could regulate various target genes ^16^.

miRNAs could also play important roles in biotic and abiotic stress ^17^. MiRNAs usually targets transcription factors and involved in adapting to stress via their positive or negative regulatory roles in plants ^12^. Thus, because of these responsive roles of miRNAs, researchers tend to investigate the role of predicted miRNAs in stress response. In this context, cowpea predicted miRNAs have been evaluated for their expression in drought tolerant and sensitive varieties ^18^.

Although, *Ae. tauschii* has been introduced as one of the most important genetic resources for the bread wheat breeding programs, but, rare studies have been focused in the detection or prediction of novel miRNAs in this species. The aim of the current study is to predict novel miRNAs in *Ae. tauschii* and their potential roles specially in stress response. We have also investigated the possibility of existence these predicted novel miRNAs in other plant species.

## Materials and methods

### Reference data sets of miRNA

To identify potential miRNAs, all previously known stem-loop and mature miRNAs of viridiplantae group were obtained (miRBase release 21.0, October 2016). 173 numbers of obtained mature miRNAs were belonged to 88 precursors of *Ae. tauschii*. Furthermore, all viridiplantae ESTs/TSAs were obtained from expressed sequence tags (ESTs) and transcriptome shotgun assembly (TSAs) databases (NCBI, October 2016). Available TSAs were belonged to assemble RNA sequences of ten representative accessions of *Ae. tauschii* that has been reported recently ^19^.

### Prediction of potential miRNAs

Blast+ 2.5.0 program downloaded from the NCBI ftp site (ftp://ftp.ncbi.nih.gov/) was used in the current study to similarity search. Known mature miRNAs were searched against ESTs/TSAs using BLASTn to separate ESTs/TSAs that could be similar with known mature miRNAs. Remained ESTs/TSAs were used for BLASTx against protein-coding sequences. Non-protein-coding sequences were used then by miReap to predict miRNA stem-loops ^20^. All predicted miRNA stem-loops were searched against known *Ae. tauschii* stem-loops using BLASTn. Some predicted miRNA stem-loops that were almost same with known *Ae. tauschii* stem-loops were eliminated. Stem-loop hairpin prediction of candidate miRNAs were folded using Mfold ^21^. An overview of our analysis methods are presented in figure 1.

**Figure 1:**
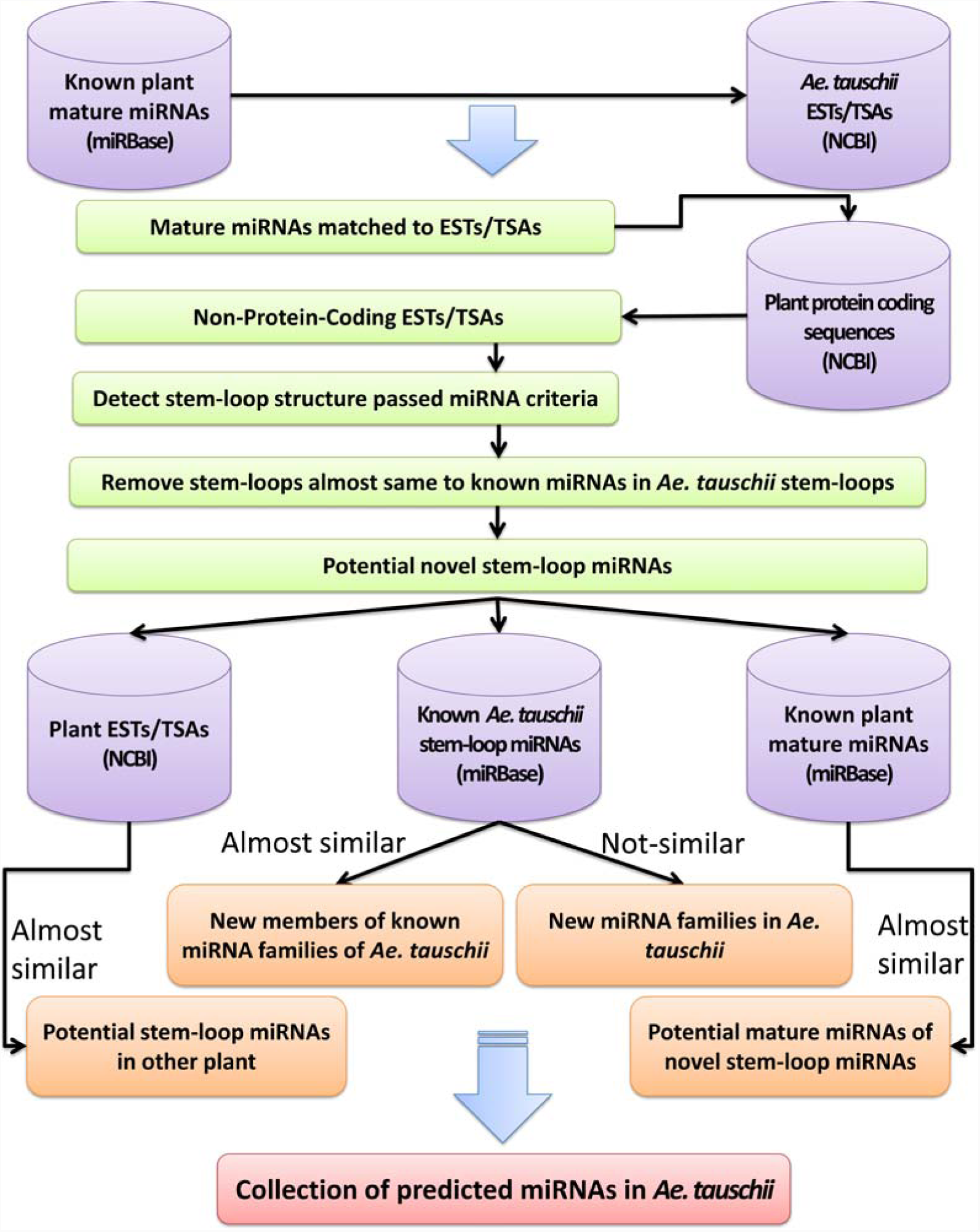
Workflow of miRNA prediction in this study.

Briefly, an identified structure was considered as a predicted stem-loop miRNA only if the following six criteria were met: (1) the sequence could fold into an appropriate stem-loop hairpin structure (2) at least one mature miRNA matched to the one arm of the stem-loop; (3) no loop between mature miRNA and its opposite sequence (4) miRNA having less than six mismatches with its opposite sequence; (5) stem-loop with higher minimal folding free energy (MFE) minimal folding free energy index (MFEI); (6) and the MFEI ranged around 0.6-0.9; ^15,16,22^. MFE was estimated using miReap and MFEI was calculated using the following equation:

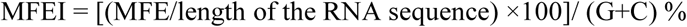

### Computational prediction of miRNA target

psRNATarget online tool ^23^ was used in this study to predict miRNA target genes. To this end, all mature miRNAs matched to the predicted stem-loops were analyzed against ESTs/TSAs of *Ae. tauschii*. Predicted targets should meet the following criteria: (1) Maximum expectation threshold = 3; (2) Length for complementarity scoring (hspsize) = 20; (3) number of top target genes for each small RNA = 200; (4) Target accessibility - allowed maximum energy to unpair the target site (UPE) = 25.0; (5) Flanking length around target site for target accessibility analysis = 17 bp in upstream and 13 bp in downstream; and (6) Range of central mismatch leading to translational inhibition = between 9 and 11 nt ^23^.

## Results

In this study using ESTs/TSAs databases nine novel miRNA stem-loops have been predicted. These stem-loops could be matched with some known mature miRNAs available in miRBase 21. Thus by aligning known mature miRNAs with predicted Stem-loops in this study we could also predict potential mature miRNAs of these novel stem-loops. Using methods mentioned before, we could predict nine novel stem-loop miRNAs including ata-miRNovel_1 to 9. MiR156, miR168, miR169, miR319, miR397 and miR530 are miRNA families that their known mature miRNAs could be matched with these novel stem-loop miRNAs. Some important features of these novel stem-loop miRNAs are presented in supplementary table 1. Furthermore, stem-loop structures of novel stem-loop miRNAs are presented in supplementary file 1. Our target prediction indicated that these novel stem-loop miRNAs could potentially regulate 136 target genes. We have observed that some important transcription factors were among these novel stem-loop miRNAs targets. In the following, predicted families of novel stem-loop miRNAs and also their characteristics are described for all novel stem-loop miRNAs.

### Novel stem-loop miRNAs almost similar to miR156 family

We have predicted two novel stem-loop miRNAs that some known mature miRNAs of miR159 family could be matched perfectly with them; including ata-miRNovel_1 and ata-miRNovel_2. These two novel stem-loop miRNAs are partially conserved (Figure 2). Our results indicated that conserved parts of these two novel stem-loop miRNAs are almost same with some known mature miRNAs. Osa-miR156c-3p, cme-miR156j and gma-miR156k are some known mature miRNAs that could be matched with ata-miRNovel_1. Likewise, bdi-miR156f-3p, stu-miR156f-5p and tae-miR156 are some other known mature miRNAs that could be matched with ata-miRNovel_2. Interestingly, known matures miRNAs that were deposited in the 5’ ends of their known stem-loops could be deposited to the 3’ end of these novel stem-loop miRNAs. It also happened vice versa. As an example, osa-miR156c-3p and bdi-miR156f-3p could be matched to the 5’ end of ata-miRNovel_1 and ata-miRNovel_2, respectively. We have also observed that known mature miRNAs of *Ae. tauschii* miR156 family (ata-miR156a,b,c,d,e-5p) could be matched to these two novel stem-loop miRNAs perfectly. However, ata-miR156a,b,c,d,e-3p and ata-miR156a,b,d,e-3p could not be matched to these two novel stem-loop miRNAs completely. These novel stem-loop miRNAs are not conserved among plant species except mature miRNAs producing regions. Our investigation among EST and TSA databases indicated that these two novel stem-loop miRNAs are similar to some *triticum* family transcripts such as GR521537.1 (EST of Perennial ryegrass etiolated seedlings) and GEDT01030793.1 (TSA of *T. polonicum*) accessions. Furthermore, our target prediction indicated that both ata-miRNovel_1 and ata-miRNovel_2 could play important role in SPL transcription factor regulation.

**Figure 2:**
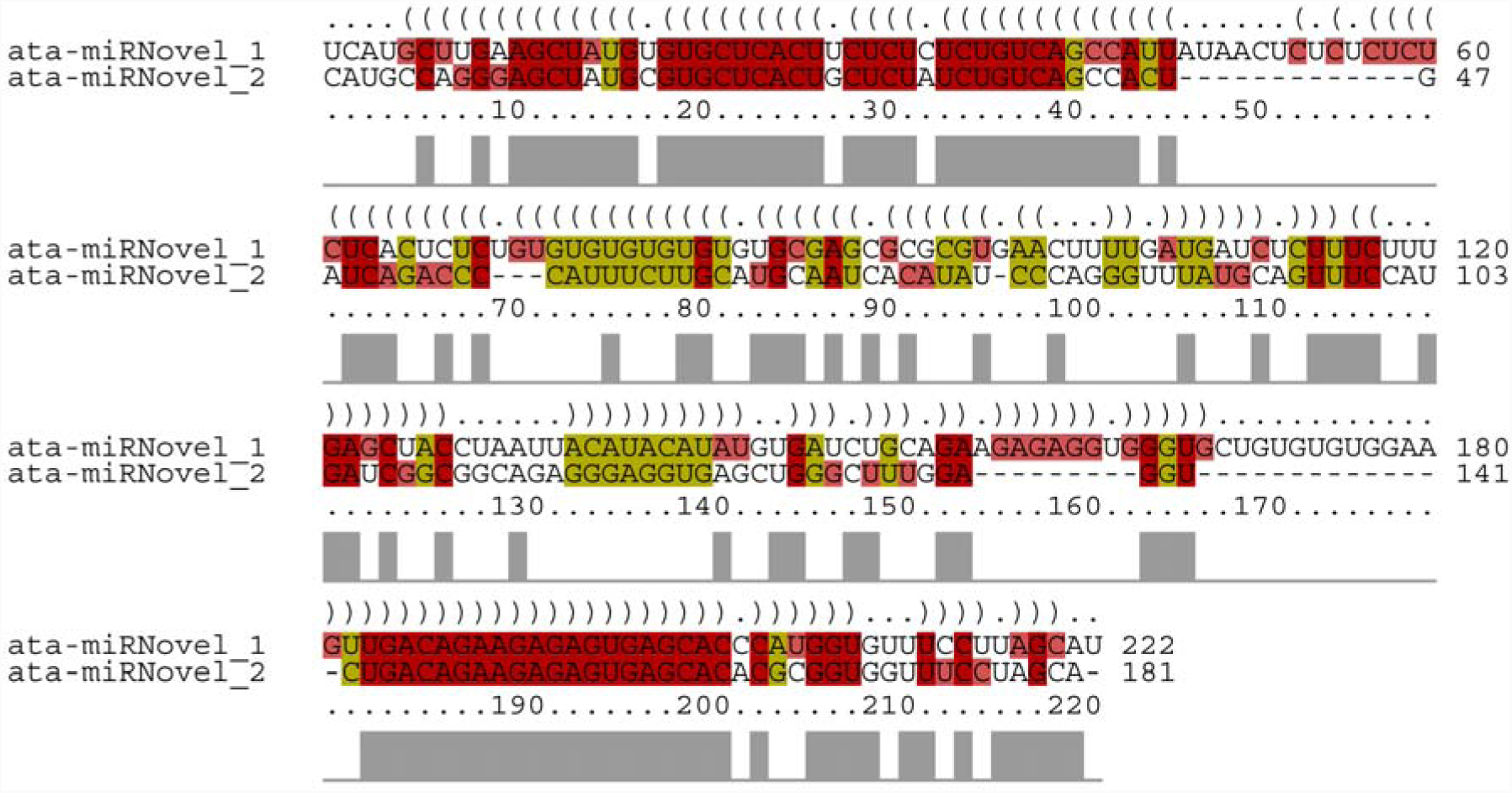
Conserved sequences between ata-miRNovel_1 and ata-miRNovel_2.

### Novel stem-loop miRNAs almost similar to miR169 family

Our analysis and predictions revealed four other novel stem-loop miRNAs belonged to miR169 family including ata-miRNovel_3, ata-miRNovel_4, ata-miRNovel_5 and ata-miRNovel_6. MFE and MFEI of these four novel stem-loop miRNAs scored −58.37 and 0.89 in average, respectively. We have observed TSAs belonged to *T. aestivum* with high similarity to some of these predicted stem-loops including HAAB01044151.1 (TSA of *T. turgidum*) and HAAB01025366.1 (TSA of *T. turgidum*) for ata-miRNovel_4 and ata-miRNovel_6, respectively. We have also observed high similarity between ata-miRNovel_3 and GEDT01056597.1 (TSA of *T. polonicum*) and also ata-miRNovel_5 and GAKM01082535.1 (TSA of *T. turgidum*). Furthermore, Blast of these novel stem-loop miRNAs against all plant known stem-loop miRNAs indicates that ata-miRNovel_3 is almost similar to bdi-miR169d. Likewise, other stem-loops including ata-miRNovel_4, ata-miRNovel_5 and ata-miRNovel_6 were almost as similar as far-miR169. Additionally, known mature miRNAs that could be matched to these stem-loops all were belonged to miR169 family. Among miR169 family in *Ae. tauschii* two members including ata-miR169c and ata-miR169i could be matched with these novel stem-loop miRNAs (Figure 3). In addition we have observed that these novel stem-loop miRNAs could regulate NF-YA transcription factors as miR169 family do. Thus, these four stem-loops could be introduced as novel stem-loop miRNAs belonged to miR169 family in *Ae. tauschii* species.

**Figure 3:**
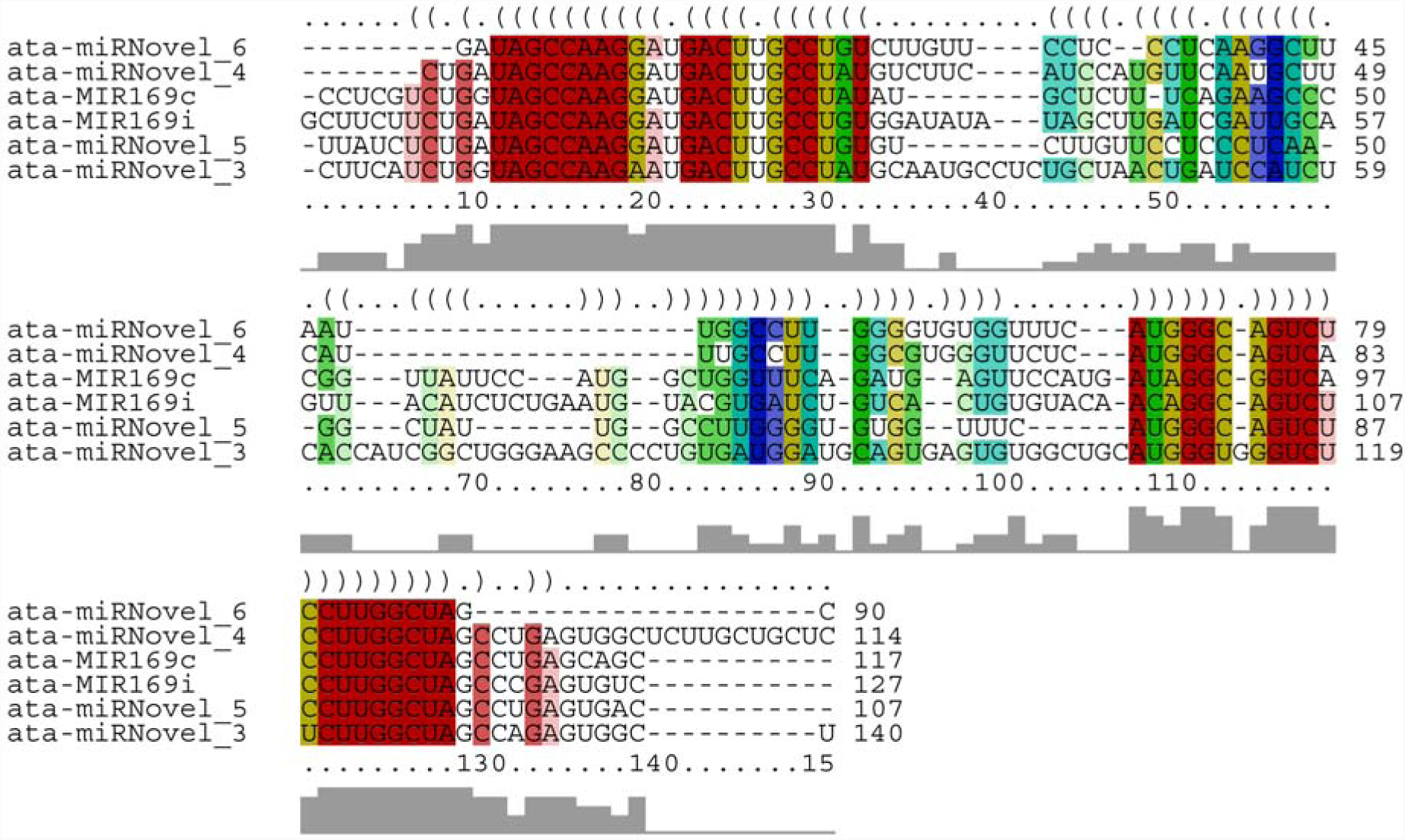
Conservation among four novel miRNAs (ata-miRNovel_3, ata-miRNovel_4, ata-miRNovel_5 and ata-miRNovel_6) with ata-miR169c and ata-miR169i.

### ata-miRNovel_7 as new Novel miRNAs almost similar to miR319 family

Our investigation indicated there is a potential stem-loop almost as similar as zma-miR319b that could be produced from *Ae. tauschii* transcripts. zma-miR319b-5p, bdi-miR319b-3p and osa-miR319b are some introduced mature miRNAs that could be matched to this novel stem-loop miRNA completely. ata-miR319-5p and ata-miR319-3p mature miRNAs of *Ae. tauschii* could be matched to this stem loop but with 2 and 4 mismatch nucleotides, respectively. MFE of this stem-loop was −104.5 and MFEI was observed as 0.43. Because of similarities between this predicted stem-loop and miR319 family, this stem-loop could be introduced as new member of this family in *Ae. tauschii*. We have observed an EST sequence (DV508145.1) with high similarity to this stem-loop sequence that was belonged to *Zea mays*. Furthermore, a TSA (GAEF01046396.1) belonged to *T. aestivum* could be matched with this novel stem-loop miRNA with high similarity. We have also observed that TCP (TEOSINTE BRANCHED/CYCLOIDEA/PCF) transcription factor could be regulated by ata-miRNovel_7 using psRNATarget. Alignment and conserved sequence areas of ata-miRNovel_7 with ata-miR319 are deposited in figure 4.

**Figure 4:**
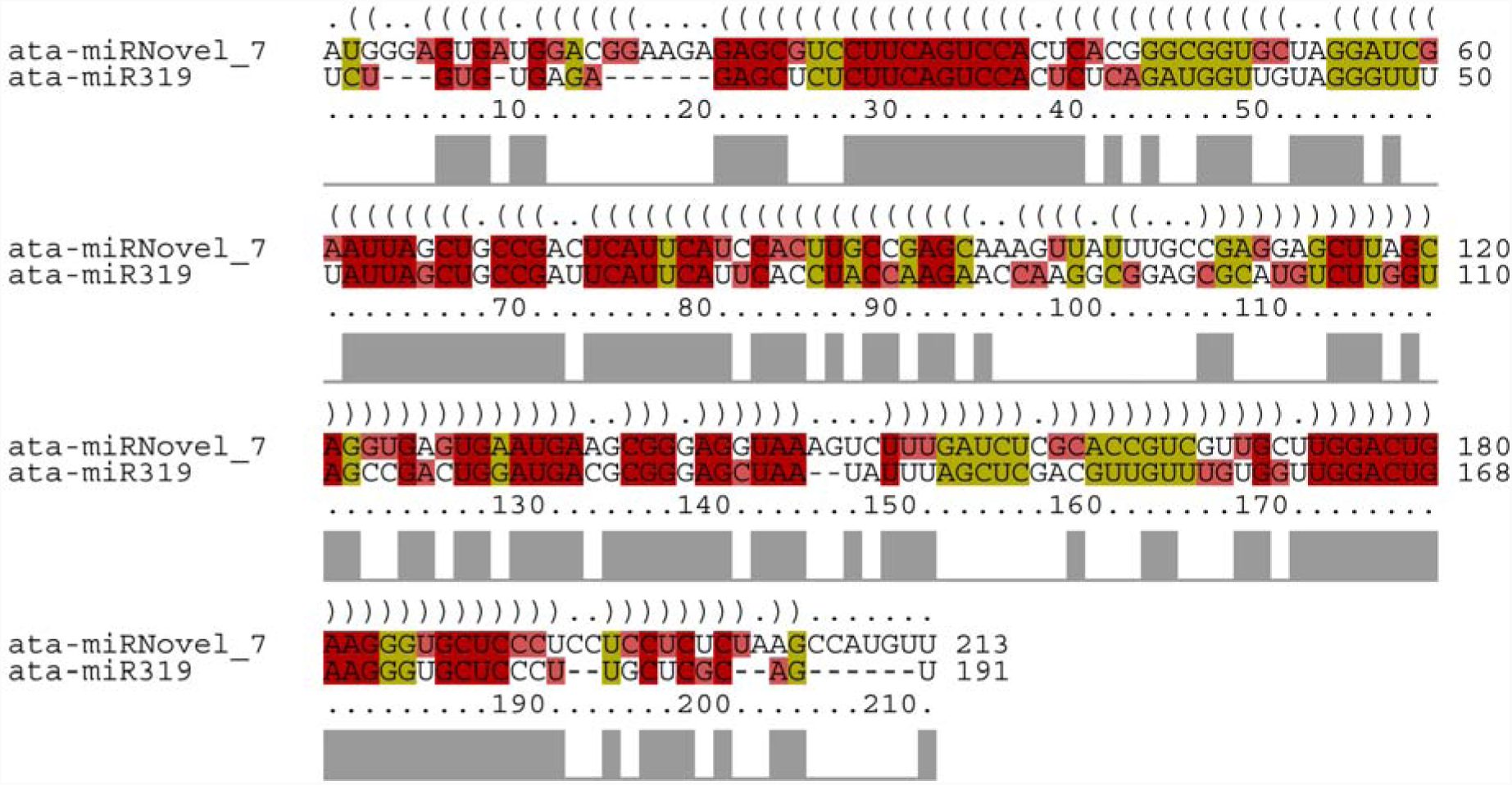
Conserved sequences between ata-miRNovel_7 with ata-miR319.

### ata-miRNovel_8 as new Novel miRNAs almost similar to miR397 family

ata-miRNovel_8 was one of the other important novel stem-loop miRNAs identified in our study. MFE and MFEI of this novel stem-loop miRNA were −56.9 and 1.15, respectively. None of *Ae. tauschii* known mature miRNAs could be matched to this novel stem-loop miRNA. However, mature miRNAs from other plant species such as ptc-miR397b, sly-miR397 and bdi-miR397b-5p could be matched to this stem-loop completely. Target prediction for predicted mature miRNA of ata-miRNovel_8 indicated that this miRNA could play important role in CBF gene regulation. Sequence of ata-miRNovel_8 was almost same with GH985139.1 (EST of *T. aestivum*) and JV942220.1 (TSA of *T. aestivum*). ata-miRNovel_8 showed the most similarity with a known stem-loop miRNA of *T. aestivum* named tae-miR397 (figure 5). Furthermore, miR397 has been not identified *Ae. tauschii* before. Thus, this predicted stem-loop could be introduced as the first member of miR397 family in *Ae. tauschii*.

**Figure 5:**
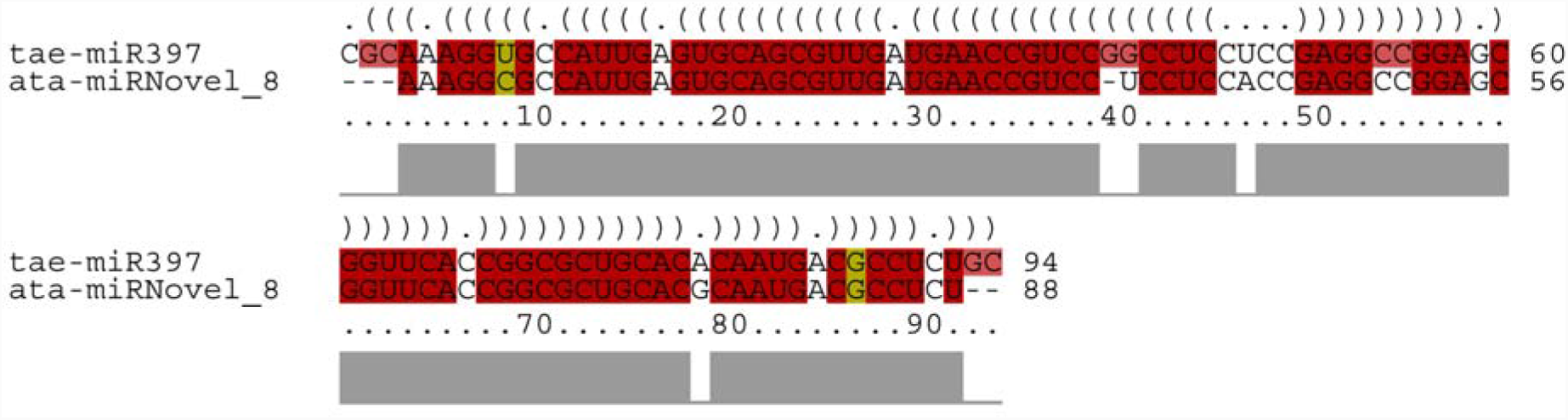
Alignment of ata-miRNovel_8 and tae-miR397.

### ata-miRNovel_9 as new Novel miRNAs almost similar to miR530 family

ata-miRNovel_9 was the last identified novel stem-loop miRNA in this study. Our results indicated that MFE and MFEI of this miRNA are −51.1 and 1.0, respectively. ata-miRNovel_9 as like as ata-miRNovel_8 could not be matched with *Ae. tauschii* known mature miRNAs. On the other hand, osa-miR530-5p and bdi-miR530a are two plant known miRNAs that could be matched with ata-miRNovel_9 completely. We have also observed that there is similarity between ata-miRNovel_9 and tae-miR530 stem-loops but not in throughout of their sequences (Figure 6). Thus, this novel stem-loop miRNA might be introduced as the first member of miR530 family in *Ae. tauschii* that has not been reported before. We have also observed high similarity between ata-miRNovel_9 and two wheat transcripts including CA655063.1 (EST of *T. aestivum*) and HP620816.1 (TSA of *T. aestivum*).

**Figure 6:**
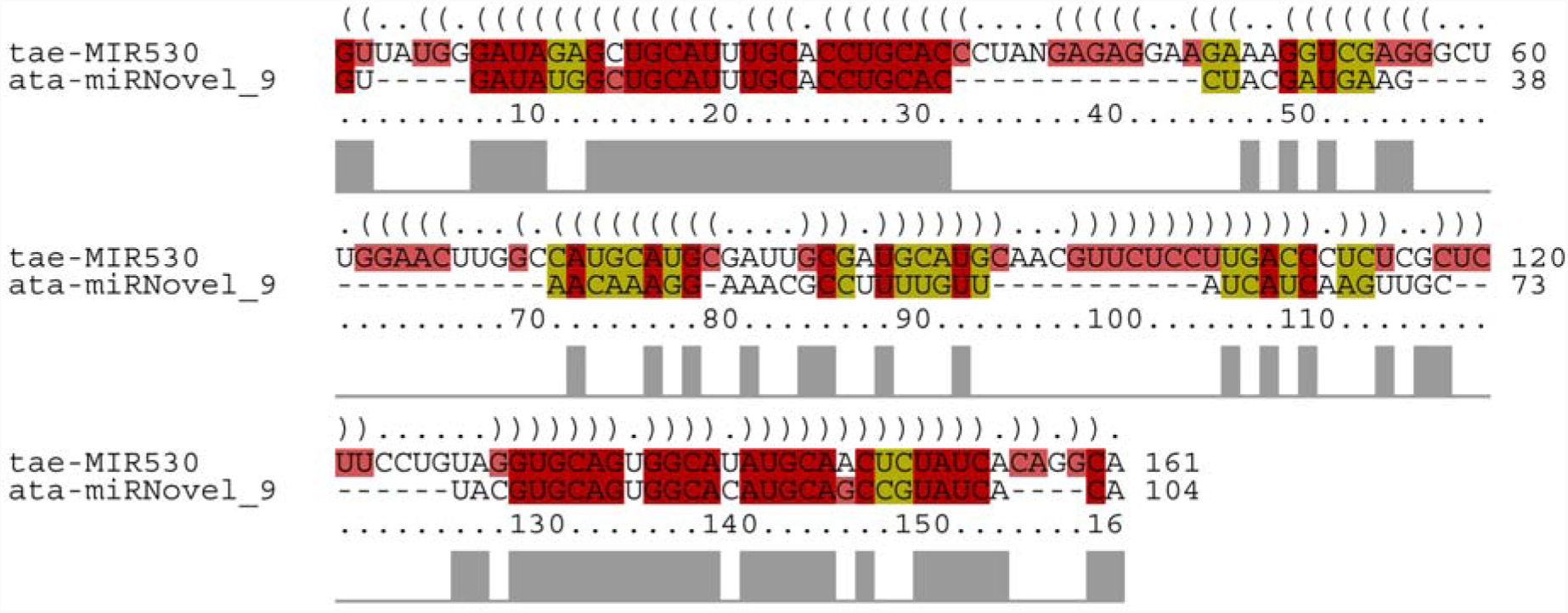
Conserved sequence between tae-miR530 and ata-miRNovel_9.

## Conclusion

In this study we have observed that each predicted stem-loop could be belonged to one of the plant known miRNA families. Ata_miRNovel_1,2 were similar to miR156 family. This family could play important role in regulation of SPL transcription factor ^24,25^. We have also observed high similarity between ata_miRNovel_3 to 6 and miR169 family. It has been shown that miR169 family are involved in regulation of NF-YA transcription factors ^26,27^. Ata_miRNovel_7 another novel identified miRNA in this study was similar to miR319 family. TCP transcription factor could be regulated by miR319 family ^28^. We have also observed similarity between ata_miRNovel_8 and miR397 family. This family could play a role in regulation of CBF gene ^29^. Previous reports emphasized the role of these families in response to biotic and abiotic stress in plants. It has been reported that miR156 could regulate tolerance to recurring environmental stress through SPL transcription factors ^30^. It has been also reported that miR156-SPLs-DFR pathway, coordinates development and abiotic stress tolerance ^31^. Furthermore, miR169 family could regulate drought tolerance through NF-YA regulation ^26,27^. In addition, TCP transcription factor as target of miR319, could control biosynthesis of the hormone jasmonic acid ^28^. MiR319 family regulates cell proliferation, leaf development and senescence through regulation of TCP transcription factor ^28,32,33^. It has been also shown that miR319 family cooperation with TCP transcription factor could play important role in response to cold, salt and drought stress ^34,35^. Additionally, the role of miR397 in cold tolerance response through CBF gene has been published before ^29^. Therefore, novel miRNAs including ata_miRNovel1 to 8 could play important roles in response to abiotic stress such as drought, salt and cold stress in *Ae. tauschii* as wild wheat relative.

In this study we have observed high similarity between sequences of novel miRNAs Stem-loop of *Ae. tauschii* and *T. aestivum* (bread wheat) transcripts. Accession numbers of these transcripts are mentioned already. According to our investigation, none of these transcripts could code a protein. Thus, these transcripts could be formed novel stem-loops in wheat that could be introduced as novel miRNAs member in Wheat. On the other hand, *Ae. tauschii* has been introduced as D genome donor of bread wheat. Therefore, it could be concluded that these wheat transcripts that are similar to new predicted stem-loops might be derived from *Ae. tauschii* through bread wheat evolution.

Overall, results of our study revealed nine novel miRNAs using ESTs/TSAs databases that showed high similarity to bread wheat transcripts. Target genes of these novel miRNAs emphasized important role of these miRNAs in response to abiotic stresses. Therefore, it could be recommended to investigate more about these novel miRNAs as parts of abiotic stress response in *Ae. tauschii* as wild wheat relative or even wheat.

## Supporting information

. Some important features of these novel stem-loop miRNAs

stem-loop structures of novel stem-loop miRNAs

