## Supplementary material for "Whole *Aegilops tauschii* transcriptome investigation revealed nine novel miRNAs involved in stress response": . Some important features of these novel stem-loop miRNAs

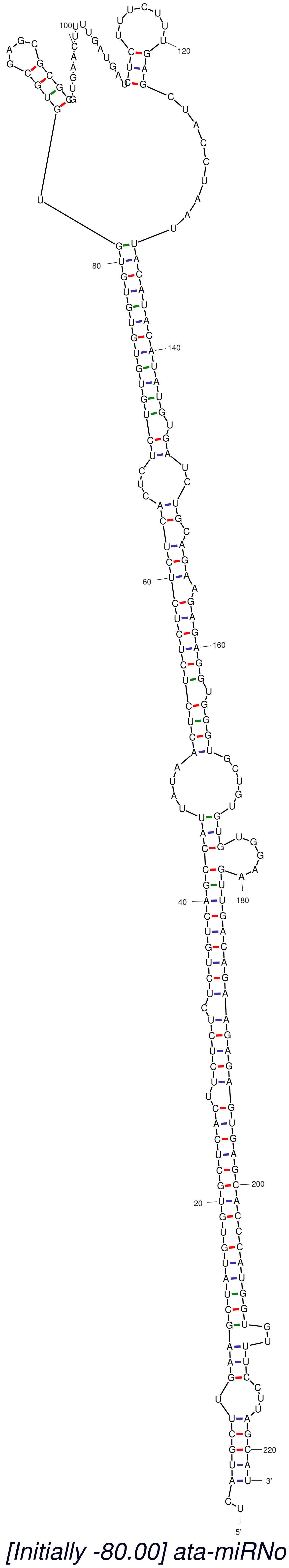


*dG = -76.09 [Initially -80.00] ata-miRNovel_1*

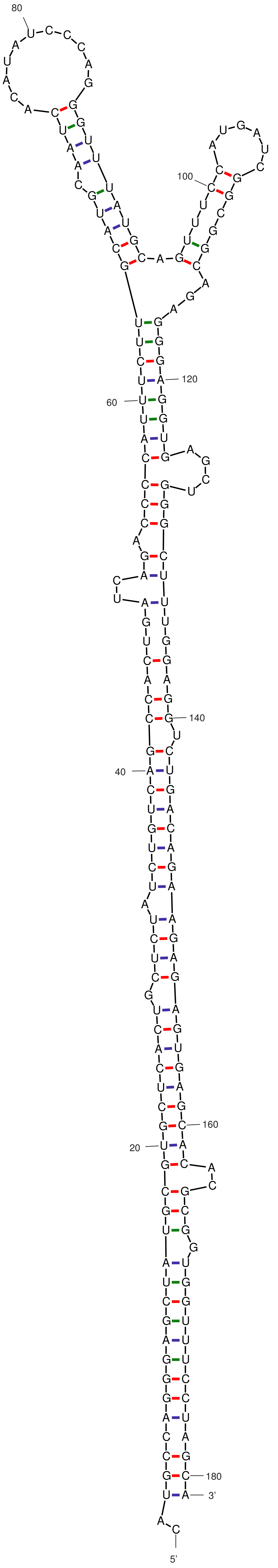


*dG = -72.20 [Initially -75.10] ata-miRNovel_2*

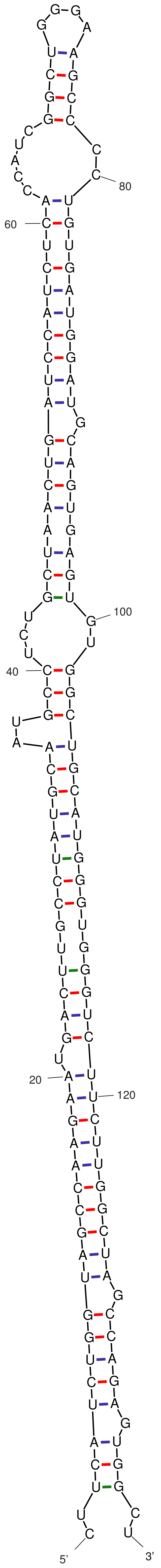


*dG = -75.90 [Initially -75.90] ata-miRNovel_3*

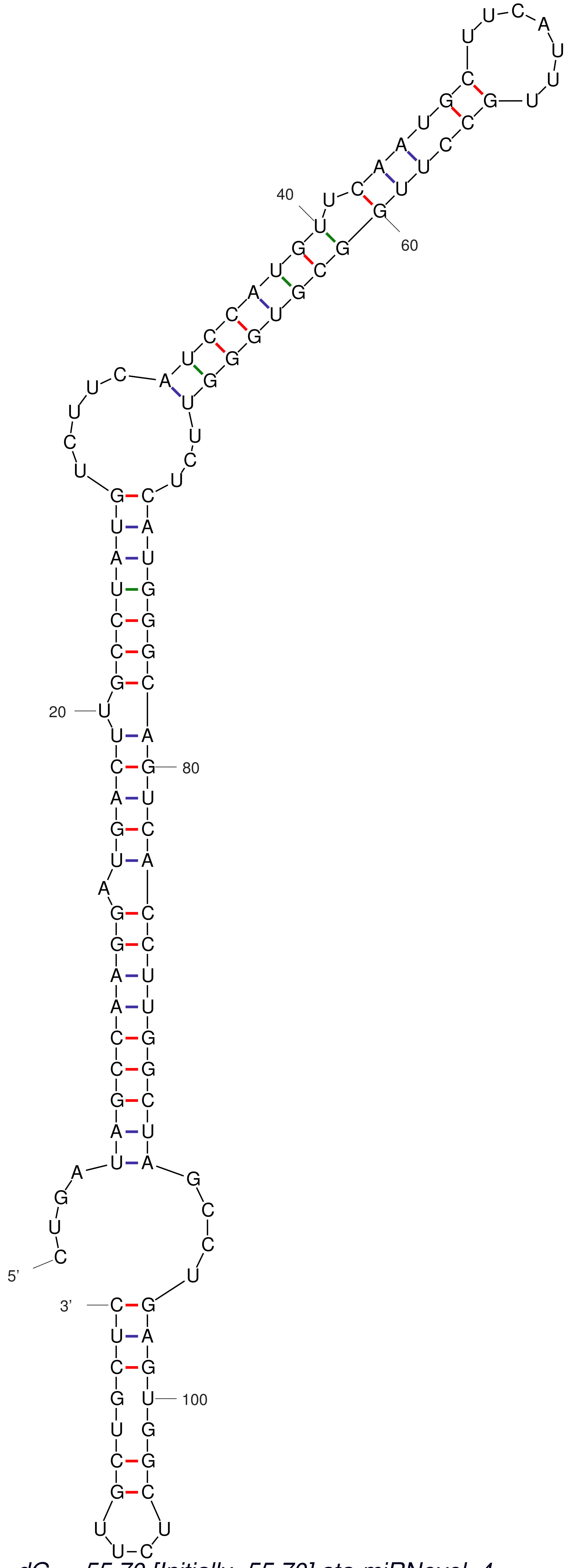


*dG = -55.70 [Initially -55.70] ata-miRNovel_4*

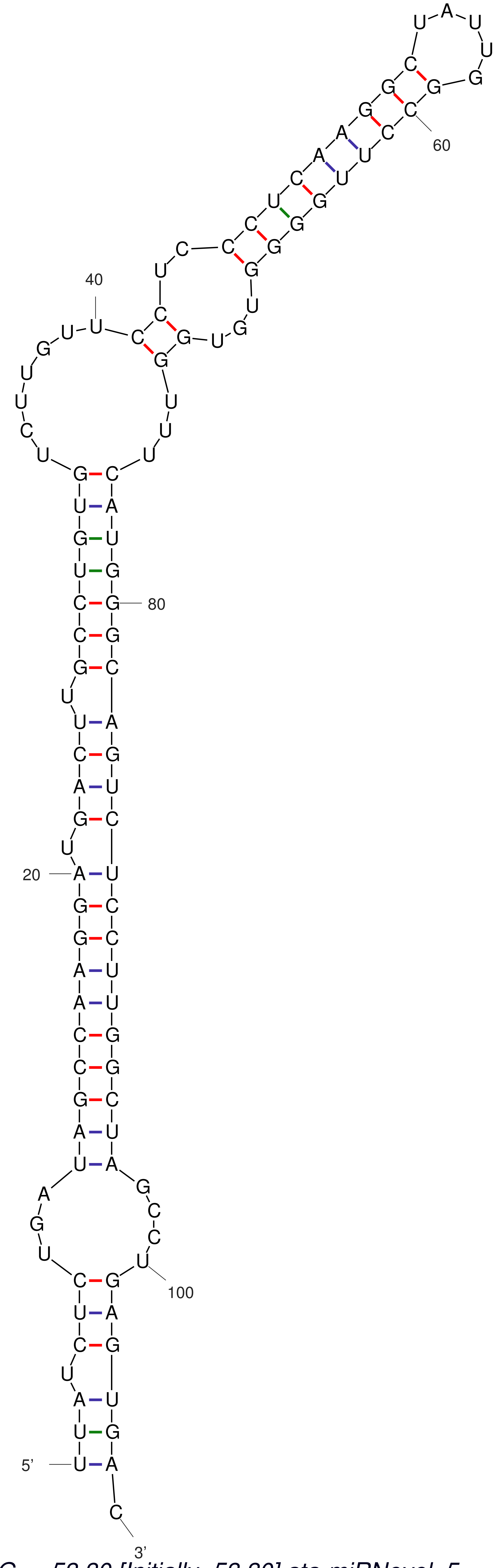


*dG = -53.30 [Initially -53.30] ata-miRNovel_5*

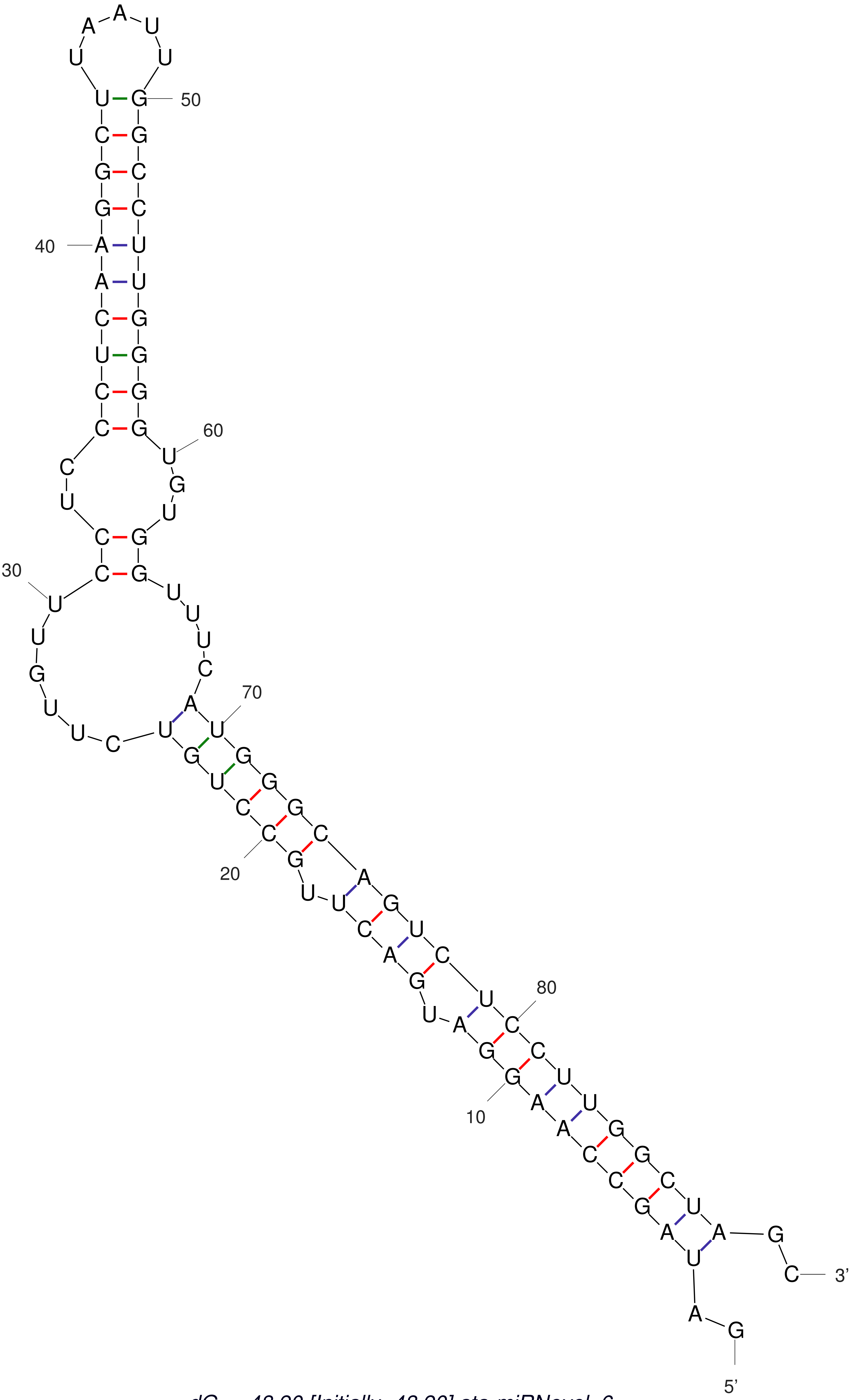


*dG = -48.90 [Initially -48.90] ata-miRNovel_6*

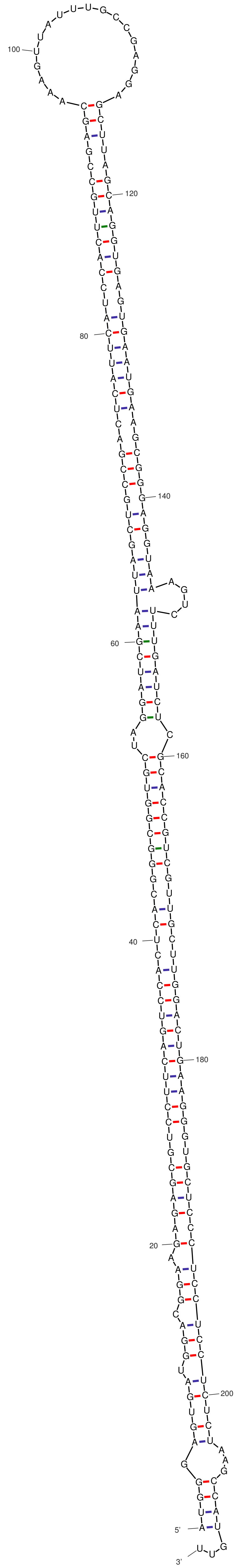

dG = -103.10 [Initially -103.10] ata-miRNovel_7

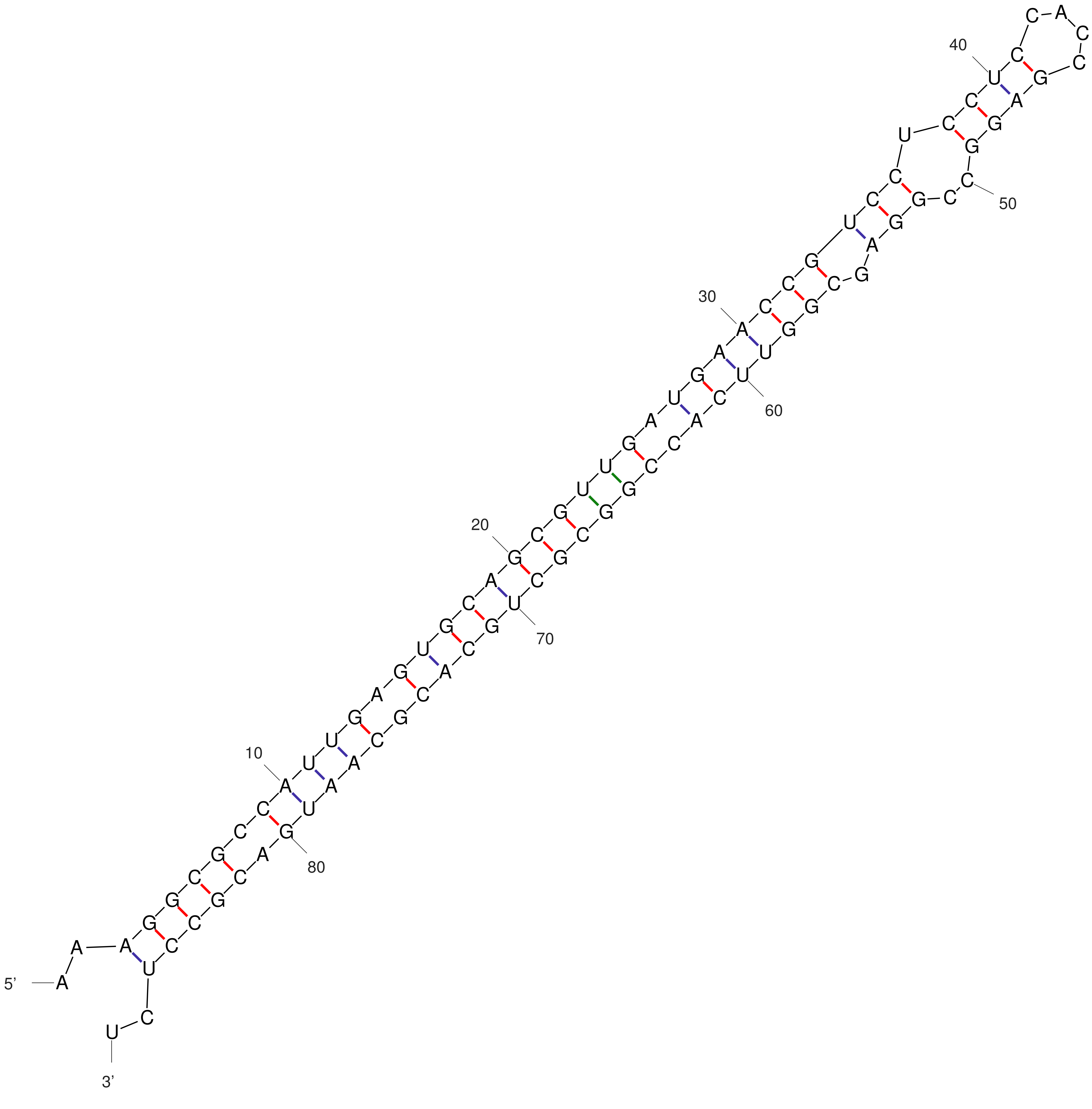


*dG = -56.30 [Initially -56.30] ata-miRNovel_8*

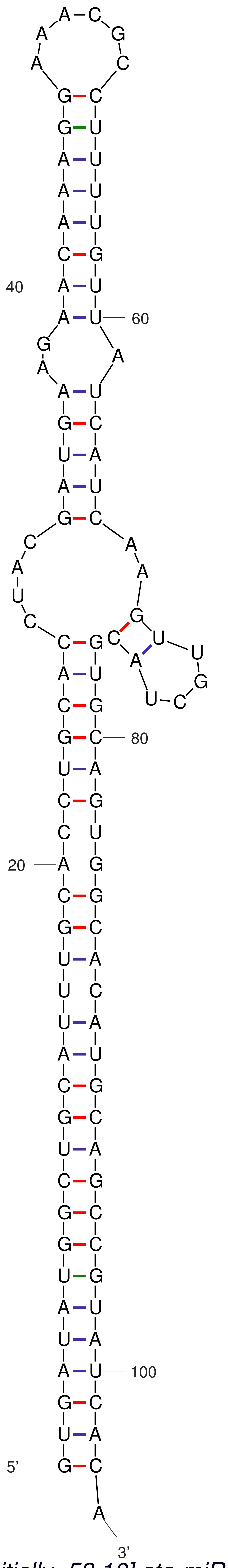


dG = -51.40 [Initially -53.10] ata-miRNovel_9
